# A versatile genetic toolbox for *Prevotella copri* enables studying polysaccharide utilization systems

**DOI:** 10.1101/2021.03.19.436125

**Authors:** Jing Li, Eric J.C. Gálvez, Lena Amend, Éva Almasi, Aida Iljazovic, Till R. Lesker, Agata A. Bielecka, Till Strowig

## Abstract

*Prevotella copri* is a prevalent inhabitant of the human gut and has been associated with plant-rich diet consumption and diverse health states. The underlying genetic basis of these associations remains enigmatic due to the lack of genetic tools. Here, we developed a novel versatile genetic toolbox for rapid and efficient genetic insertion and allelic exchange applicable to *P. copri* strains from multiple clades. Enabled by the genetic platform, we systematically investigated the specificity of polysaccharide utilization loci (PULs), and identified four highly conserved PULs for utilizing arabinan, pectic galactan, arabinoxylan and inulin, respectively. Further genetic and functional analysis of arabinan utilization systems illustrate that *P. copri* has evolved two distinct types of arabinan-processing PULs (PUL^Ara^) and that the type-II PUL^Ara^ is significantly enriched in individuals consuming a vegan diet compared to other diets. In summary, this genetic toolbox will enable functional genetic studies for *P. copri* in the future.

## Introduction

The complex microbial communities residing in the intestine affect the physiology of the host influencing the balance between health and disease (Bäckhed et al., 2005; Hooper, 2009). Yet, extensive interpersonal variability in the human gut microbiota composition and function complicates the establishment of links between the presence of specific bacterial gene content to human phenotypes. Functional genetic study of these bacteria is essential to dissect the genetic basis underlying the microbe-driven host phenotypes. However, many commensal bacterial species have so far eluded efforts for genetic engineering.

For instance, no genetic tools have been established for *Prevotella copri*, a common human gut microbe, whose prevalence and relative abundance have been linked to various beneficial and detrimental effects on human health (Claus, 2019; Ley, 2016; Maeda and Takeda, 2019). Specifically, *P. copri* has been found to be enriched in individuals at risk for rheumatoid arthritis (Alpizar-Rodriguez et al., 2019; Scher et al., 2013; Wells et al., 2020) and in patients with enhanced insulin resistance and glucose intolerance (Pedersen et al., 2016). Conversely, others found *P. copri* to be positively correlated with improved glucose and insulin tolerance during intake of fiber-rich prebiotic diets (Kovatcheva-Datchary et al., 2015; De Vadder et al., 2016). Besides the lack of tools for genetic engineering, the establishment of functional links between *P. copri* and disease outcomes has been additionally complicated by its fastidious nature *in vitro*, tremendous strain-level diversity, resulting in the recent recognition of multiple genetically distinct clades and the lack of corresponding diverse isolates (Tett et al., 2019).

In contrast to *P. copri*, members of the genus *Bacteroides*, as the best example, have been extensively studied via a variety of genetic tools (Bencivenga-Barry et al., 2020; García-Bayona and Comstock, 2019; Goodman et al., 2011; Koropatkin et al., 2008; Lim et al., 2017; Mimee et al., 2015). These studies have, for instance, identified genes required for various bacterial physiological functions and provide approaches to investigate bacteria-host interactions. Of those, the genes for degrading plant- and animal-derived polysaccharides that are resistant to human digestion have been highlighted due to their important role in affecting bacterial fitness in the microbiome (Kaoutari et al., 2013; Porter and Martens, 2017; Wexler and Goodman, 2017). These genes are typically organized in so called polysaccharide utilization loci (PULs) that differ in polysaccharide specificity (Koropatkin et al., 2012). PULs are defined by the presence of one or more genes homologous to *Bacteroides thetaiotaomicron susD* and *susC* encoding outer membrane proteins that bind and import starch oligosaccharides (Martens et al., 2009; Shipman et al., 2000). The SusC/D protein complex (Glenwright et al., 2017) cooperates with diverse carbohydrate-degrading enzymes (CAZyme), e.g., glycosyl hydrolases (GHs) and polysaccharide lyases (PLs), which are typically encoded in close proximity to the *susC/D* homologs in the genome. Most PULs in *B. thetaiotaomicron* contain genes encoding sensor-regulator systems, such as hybrid two-component systems (HTCSs) (Sonnenburg et al., 2006; Xu et al., 2003). HTCS proteins are chimeric proteins harboring the functional domains of a periplasmic sensor, a histidine kinase, and a DNA-binding response regulator enabling HTCSs to recognize distinct signal components degraded from complex carbohydrates and to initiate the upregulation of CAZyme-encoding genes in a positive feedback loop (Sonnenburg et al., 2006, 2010).

Notably, higher prevalence of intestinal *Prevotella* spp. was found in populations consuming a plant-rich diet, e.g., vegetarians in Western populations (De Filippo et al., 2010; Fragiadakis et al., 2019; Ruengsomwong et al., 2016; Wu et al., 2011) suggesting that they encode efficient machineries for degradation of plant-derived polysaccharides. Yet, due to the lack of genetic tools, the characterization of carbohydrate utilization has been limited to bioinformatic and phenotypic studies (Fehlner-Peach et al., 2019; De Filippis et al., 2019). Specifically, two recent studies described extensive variability among clades and also strains within clades in the ability to directly utilize diverse complex plant carbohydrates (Fehlner-Peach et al., 2019; Tett et al., 2019). While combinations of comparative genomics and phenotypic assays can be used to predict the ability to utilize specific polysaccharides, such approaches rely on well-characterized genetic elements as a reference, which makes it difficult to identify genes with unknown functions and thereby hinders the establishment of casual relationship between the genetic content and phenotypes. Moreover, substrate predictions based on gene annotations from genetically distinct bacteria might be incomplete or inaccurate. This observation is supported by the presented data below that some *P. copri* strains harboring PULs lacking marker genes can still grow on the substrates for those PULs suggesting the functional redundancy between various PUL components.

Here, we described a newly established genetic toolbox for approaching gene insertion, deletion, and complementation in *P. copri*. Using the genetic tools as well as high-throughput sequencing and bioinformatic analysis, we identified four highly conserved *P. copri* PULs responsible for utilization of specific plant polysaccharides via HTCS activation, and demonstrate that *P. copri* species have evolved two types of arabinan processing PULs. These studies not only build up a universal genetic manipulation system for an abundant bacterial species in the microbiome, but also present its applications on future efforts of understanding *P. copri* biology, e.g. nutrient acquisition. Because the workflow of establishing the genetic manipulation system for *P. copri* can be potentially modified and applied to other underexplored gut bacteria, our studies shed light on the future microbiome research on intricate interactions between bacteria-bacteria and host-bacteria during human health and disease.

## Results

### Development of conjugation-based gene insertion system for *P. copri*

As targeted gene inactivation approaches enable gene function studies, they are frequently carried out in *Bacteroides* spp. by transferring a suicide plasmid from a donor strain into the recipient followed by selection of bacterial clones which underwent homologous recombination (Bencivenga-Barry et al., 2020; García-Bayona and Comstock, 2019; Koropatkin et al., 2008). To adapt the system for *P. copri*, we considered several key differences between *Bacteroides* spp. and *P. copri* including oxygen and antibiotic sensitivity as well as promoter sequences driving expression of the selectable marker gene.

Because oxygen exposure has been reported to promote mating between *Escherichia coli* (donor) and *Bacteroides* spp. (recipient) (Salyers et al., 1999), conjugation for *Bacteroides* spp. is routinely performed for at least 15 hours under aerobic conditions followed by transferring the cultures to anaerobic conditions permitting growth (Bencivenga-Barry et al., 2020; García-Bayona and Comstock, 2019). We initially tested aerotolerance of three *P. copri* strains, the type strain (DSM18205) and two strains (HDD04 and HDB01) from our lab collection containing recent isolates from healthy and diseased individuals (Figure 1A and Table S1). Exposure to air decreased viability three to four orders of magnitude for the *P. copri* strains within only four hours, which is in sharp contrast to *B. thetaiotaomicron* that displayed only a 28.6% drop in viability (Figure S1A). Hence, all genetic manipulations for *P. copri* were subsequently carried out in anaerobic conditions.

**Figure 1.**
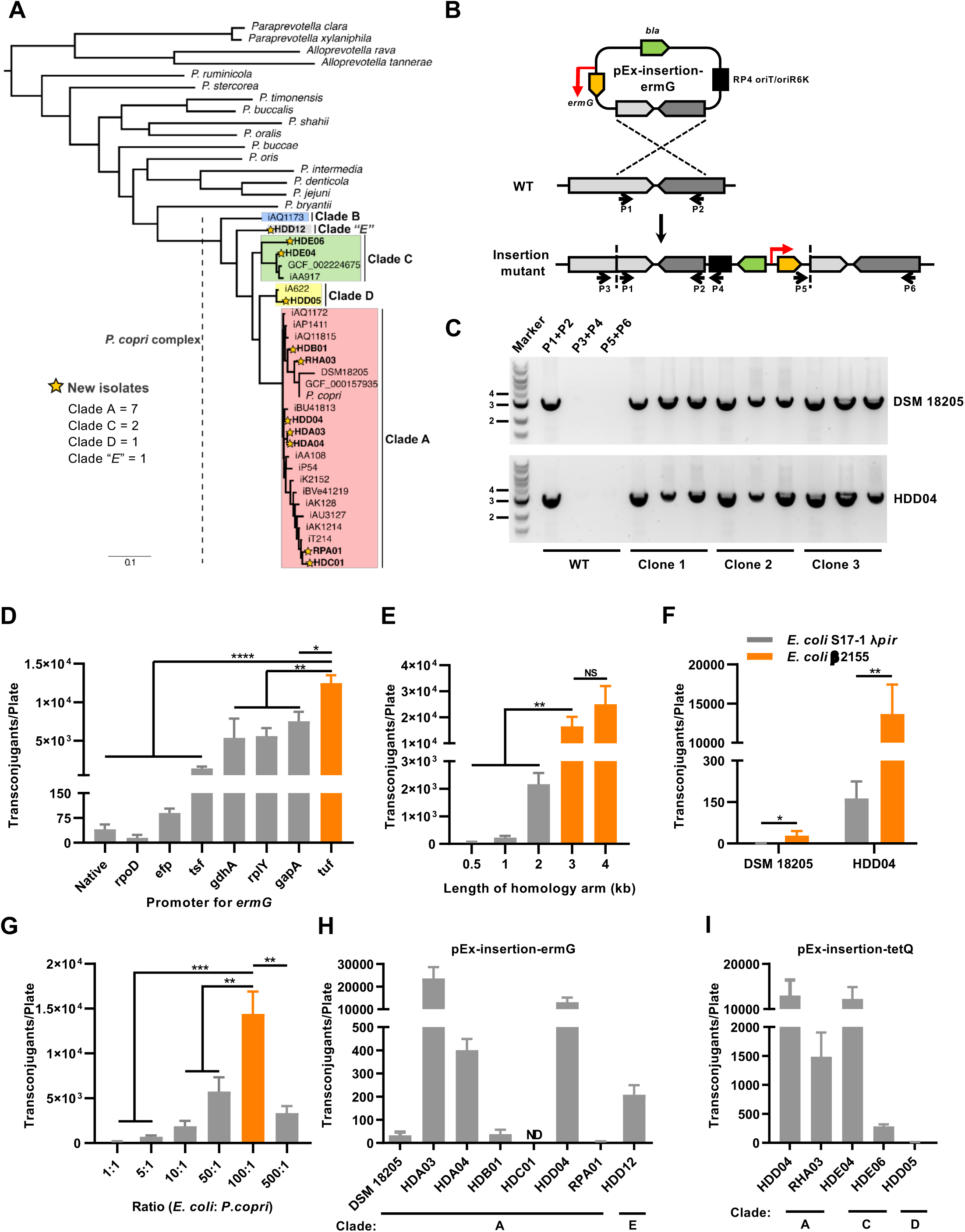
Development of a conjugation-based gene insertion platform for *P. copri* strains from multiple clades. (A) Phylogenetic tree of *P. copri* species complex using reference strains (n=17) from four *P. copri* clades (Tett et al., 2019) and novel isolates (n=11) used in this study. *P. copri* clades are indicated by different colors. (B) Schematic illustration for targeted gene insertion system in *P. copri*. Primer binding sites (P1-P6) are indicated. (C) Detection of plasmid integration in *P. copri* DSM 18205 and HDD04 by PCR. (D-H) Optimization of conjugation efficacy: Influence on yield of transconjugants of promoter sequences of selection marker (D), length of homology arm (E), donor *E. coli* strain (F), conjugation ratio of donor to recipient strains (G), recipient *P. copri* strains that are erythromycin-sensitive (H) or tetracycline-sensitive (I). Comparisons in (D-G) were performed using *P. copri* HDD04. Values and error bars represent the mean of at least three biological replicates and their standard deviations (SDs), respectively. Statistical significance between groups was calculated by Student’s *t* test (*p < 0.05; **p < 0.01; ***p < 0.001; and ****p < 0.0001; NS, p > 0.05, not statically significant; ND, not detectable)

*E. coli* S17-1 λpir is commonly used as a donor for *Bacteroides* spp. in aerobic conditions, but it may show impaired growth under anaerobic conditions. Hence, we compared anaerobic growth of *E. coli* S17-1 λpir and another donor strain, *E. coli* β2155 (Dehio and Meyer, 1997; Demarre et al., 2005). Notably, while *E. coli* β2155 is auxotrophic for diaminopimelic acid (DAP), it displayed in the presence of DAP faster and more robust anaerobic growth compared to *E. coli* S17-1 λpir both in liquid culture and on agar plates (Figure S1B and S1C). Thus, we further tested the possibility of using *E. coli* β2155 as the donor for delivering vectors into recipient *P. copri*.

The suicide plasmid, pExchange-tdk, is extensively used for gene deletion in *Bacteroides* spp. (Koropatkin et al., 2008). This plasmid possesses (1) a R6K origin limiting it to replicate only in host bacteria carrying the *pir* gene, and (2) an erythromycin resistant gene (*ermG*) for selecting *Bacteroides* transconjugants. Firstly, no *pir* homologs were identified in *P. copri* strains, suggesting the feasibility of using pExchange-tdk based vectors for plasmid integration in *P. copri*. Secondly, the susceptibility of *P. copri* strains to erythromycin was tested. From one type strain and 11 distinct strains of our collection representing four *P. copri* clades, eight strains from two clades (Figure 1A) were sensitive to erythromycin driving us to initially utilize an erythromycin-based selection system (Table S2). To achieve a stable expression of *ermG* in *P. copri*, we inserted a strong promoter of a *P. copri* housekeeping gene, i.e. elongation factor Tu gene (*tuf*) (Figure S1D), in front of the *ermG* coding sequence into pExchange-tdk and removed the counterselection marker (thymidine kinase gene). These modifications resulted in a new shuttle vector referred to as pEx-insertion-ermG (Figure 1B and S2A). To consider potential negative positional effects for *P. copri* growth due to plasmid insertion, we individually cloned three different 3-kb regions (DSM18205_00642-43, 00941-42, and 02334-35) containing the 3’-end coding sequences of genes from *P. copri* DSM 18205, as homology arms for approaching plasmid integration without disrupting any functional genes. In addition, these DNA regions are relatively conserved in genomes of our *P. copri* isolates, allowing us to rapidly expand the testing into different strains.

The initial conjugation was performed between *E. coli* β2155 carrying the respective plasmids and *P. copri* DSM 18205. Of note, *E. coli* β2155 can grow weakly on BHI blood agar plates even without DAP based on our observations. Hence, after the co-incubation of *E. coli* and *P. copri* on BHI blood agar with DAP, besides erythromycin, we additionally added gentamicin to inhibit the *E. coli* donor and positively select for *P. copri* transconjugants. This selection yielded eight erythromycin resistant (Erm^R^) colonies only when pEx-insertion-ermG-DSM18205_02334-35 was used, which indeed suggests the presence of positional effects (Figure 1B). Since DSM18205_02334 encodes a putative β-glycosidase gene (*bgl*), we refer to the plasmid as pEx-insertion-ermG-DSM-bgl. Integration of the transferred plasmid was determined in three individual colonies and colony PCR amplified fragments expected for successful integration (Figure 1C and Table S1). DNA sequences of PCR products were further confirmed by Sanger sequencing (data not shown). We next attempted conjugation using pEx-insertion-ermG-HDD04-bgl and HDD04 as the conjugation recipient, which strikingly resulted in a more than 400-fold higher number of Erm^R^ colonies (Figure 1H) suggesting a large strain variability in conjugation efficiency and prompting us to further optimize the approach.

### Optimization of conjugation-based gene insertion for *P. copri*

Building on these proof-of-concept data, the genetic elements of the plasmid and the conjugation procedures were systematically adjusted by varying factors that likely affect the conjugation efficiency. This included the promoter of *ermG*, the length of homology arm for plasmid integration, the donor *E. coli* strains and ratio of donor to recipient for conjugation, as well as the recipient *P. copri* strains.

First, an efficient expression of the selection marker is the prerequisite to obtain Erm^R^ transconjugants. Besides the promoter of the *tuf* gene, we chose another six different promoters of housekeeping genes showing diverse gene expression in *P. copri* HDD04 in BHI liquid media supplemented with fetal bovine serum (BHI+S) (Figure 1D and S1D; Table S5). Notably, the numbers of transconjugants varied approximately 300-fold depending on the promoter, but none of the other promoters yielded higher numbers than *tuf* suggesting that high *ermG* expression levels are required for transconjugant survival under erythromycin selection (Figure 1D and S1D). Second, comparison of homology arms between 0.5-kb to 4-kb demonstrated the highest yield of transconjugants with homology arms of 3- and 4-kb (Figure 1E). Third, we assessed the above-mentioned *E. coli* strains as conjugation donors. In line with the ability for anaerobic growth, conjugation with donor *E. coli* β2155 increased the yield of transconjugants for both DSM 18205 and HDD04 approximately 85-fold compared to *E. coli* S17-1 λpir, indicating the advantage of donor strain for advancing anaerobic conjugation (Figure 1F). More transconjugants were also obtained as ratio of donor to recipient increased until 100:1, after which it decreased again (Figure 1G). Last, we evaluated a larger panel of Erm^S^ *P. copri* strains using strain-specific homology arms as extensive sequence variations among strains are present. Except for one strain with undetectable production of Erm^R^ colonies all other seven strains exhibited extensive diversity in the number of transconjugants varying by approximately 10^4^-fold between the strains with the lowest (RPA01, mean=5 CFUs) and highest (HDA03, mean=2.4×10^4^ CFUs) yield (Figure 1H).

While these iterative improvements allowed the targeted insertion in seven strains from the clade A and “E” (a newly observed clade, unpublished observation), the other four strains from the clade A, C and D could not be assessed due to their Erm^R^ phenotype (Table S2). Hence, we screened the antibiotic susceptibility of HDD04 and DSM 18205 identifying tetracycline, chloramphenicol, and spectinomycin as additional selective antibiotics (Table S2). Next, *P. copri* HDD04 was utilized to test the feasibility of tetracycline, chloramphenicol, and spectinomycin resistance genes, i.e. *tetQ*, *catA*, *aadA*, for selection. Conjugation using pEx-insertion carrying *tetQ* (pEx-insertion-tetQ-bgl, Figure S2B), but not *catA* or *aadA*, successfully resulted in HDD04 transconjugants after selection with the respective antibiotics. We therefore performed the conjugation for four Erm^R^ but tetracyclin-sensitive (Tet^S^) strains, i.e., RHA03, HDE04, HDE06, and HDD05, followed by tetracycline selection, resulting in tetracycline resistant (Tet^R^) transconjugants (mean=8.3 to 1.2×10^4^ CFUs) (Figure 1I).

In summary, the comprehensive and stepwise adaptation of a gene insertion system to a genetically inaccessible bacterium, i.e. *P. copri,* illustrates the significant influence of multiple variables for a successful production of genetic mutants, thereby providing a valuable template for initiating the construction of genetic tools for other commensals in the future. These experiments together demonstrate the feasibility of gene insertion in *P. copri* strains from distinct clades enabling functional studies.

### Genetic inactivation of PUL regulators to identify their polysaccharide substrates

The prevalence and relative abundance of *P. copri* has been linked to plant-rich diets in humans and mouse models (De Filippo et al., 2010; Fragiadakis et al., 2019; Gálvez et al., 2020; Kovatcheva-Datchary et al., 2019; Ruengsomwong et al., 2016; Wu et al., 2011), yet the underlying molecular mechanism is still poorly understood. To demonstrate the utility of our gene insertion system in identifying gene functions, we decided to investigate the genetic basis for utilization of distinct carbohydrates in *P. copri*. Since no available chemically-defined cultivation system for *P. copri* exist thus far, we modified the minimal medium (MM) (Martens et al., 2008) originally used for cultivation of *B. thetaiotaomicron* by supplementing additional defined nutrients (see Methods), enabling the growth of all *P. copri* strains tested from our strain collection (n=12) with glucose as a sole carbon source (Table S3). Specifically, 10 out of 12 strains reached a maximal optical density (OD_600_ max) of 0.6-1.0 in MM+Glucose overnight, while two strains (HDA03 and RHA03) showed only moderate growth (approximately OD_600_ max 0.3). This minimal medium was then used to extensively characterize polysaccharide utilization in HDD04, as it showed robust growth in MM and high number of transconjugants. HDD04 grew on various plant cell wall pectins, such as arabinan, arabinogalactan, and arabinoxylan (Table S3). Beyond the utilization of plant cell wall glycans, HDD04 also showed growth on plant and animal cell storage carbohydrates such as inulin (0.651±0.016) and glycogen (0.818±0.026), but grew poorly on levan (0.122±0.009), and could not grow on starch. In parallel, PULs were identified in *P. copri* using a bioinformatic approach PULpy (Stewart et al., 2018) followed by manual curation. Specifically, the PUL repertoire of HDD04 was predicted based on *susC/D*-like pairs resulting in 29 PULs in comparison with 19 PULs in the *P. copri* reference strain (DSM 18205) (Table S3), suggesting a much broader carbohydrate utilization capability of HDD04 compared to DSM 18205. CAZymes surrounding the *susC/D*-like pairs were annotated using a bioinformatic approach (dbCAN2 tool) (Zhang et al., 2018).

To directly link distinct PULs and growth phenotypes on polysaccharides, we focused on HTCS genes, the typical activator associated with PULs (Sonnenburg et al., 2006, 2010). Genome-wide screening for HTCS genes in HDD04 by homology search using the known domains of HTCS (Terrapon et al., 2015) combined with a protein BLAST on National Center for Biotechnology Information (NCBI) identified ten gene candidates as our targets. We associated nine out of ten HTCS gene candidates with their closest predicted PULs (e.g. HDD04_00018 is named as *htcs*-PUL3) with only HDD04_0019 being a solitary HTCS gene.

Ten HTCS insertion mutants were generated by integrating a modified pEx-insertion-ermG plasmid (Figure S2C) into the coding sequences of their periplasmic sensor domains followed by a screening for growth defects in MM plus polysaccharides to identify their respective substrates (Figure 2A). Of note, to block potential effects of transcriptional and translational readthrough for the HTCS genes after plasmid integration, pEx-insertion-ermG was modified to include T1-T2 terminators and TAA encoding stop codon in front of the cloned homology arm, respectively (pEx-insertion-ermG-T1T2; Figure 2A and Figure S2C). A strain with plasmid integrated into an intergenic region (between HDD04_00165 and 00166) was utilized as a positive control. The polysaccharides (n=15) that can support the growth of HDD04 to an OD_600_ max of > 0.2 within 120 hours were investigated (Figure 2B). Compared to the control strain, 6/10 HTCS gene mutants displayed similar growth patterns, e.g., *htcs*-PUL3, as shown in Figure 2B, demonstrating that they are not essential for growth on the tested polysaccharides. Strikingly, the other four HTCS mutants each showed dramatic growth defects (OD_600_ max < 0.1) on only one specific polysaccharide (Figure 2C and 2D). Specifically, gene disruptions of *htcs*-PUL14, -PUL21, - PUL24, and -PUL26 abolished the capacities of HDD04 grown on arabinan, pectic galactan, arabinoxylan, and inulin, respectively (Figure 2C and 2D). To link the HTCS gene and the nearest PUL for easy identification, we have tentatively designated these genes as *htcs*^D_Ara^ (*htcs*-PUL14, HDD04_02372), *htcs*^D_PecGal^ (*htcs*-PUL21, HDD04_02939), *htcs*^D_AraXyl^ (*htcs*-PUL24, HDD04_03129), and *htcs*^D_Inu^ (*htcs*-PUL26, HDD04_03217).

**Figure 2.**
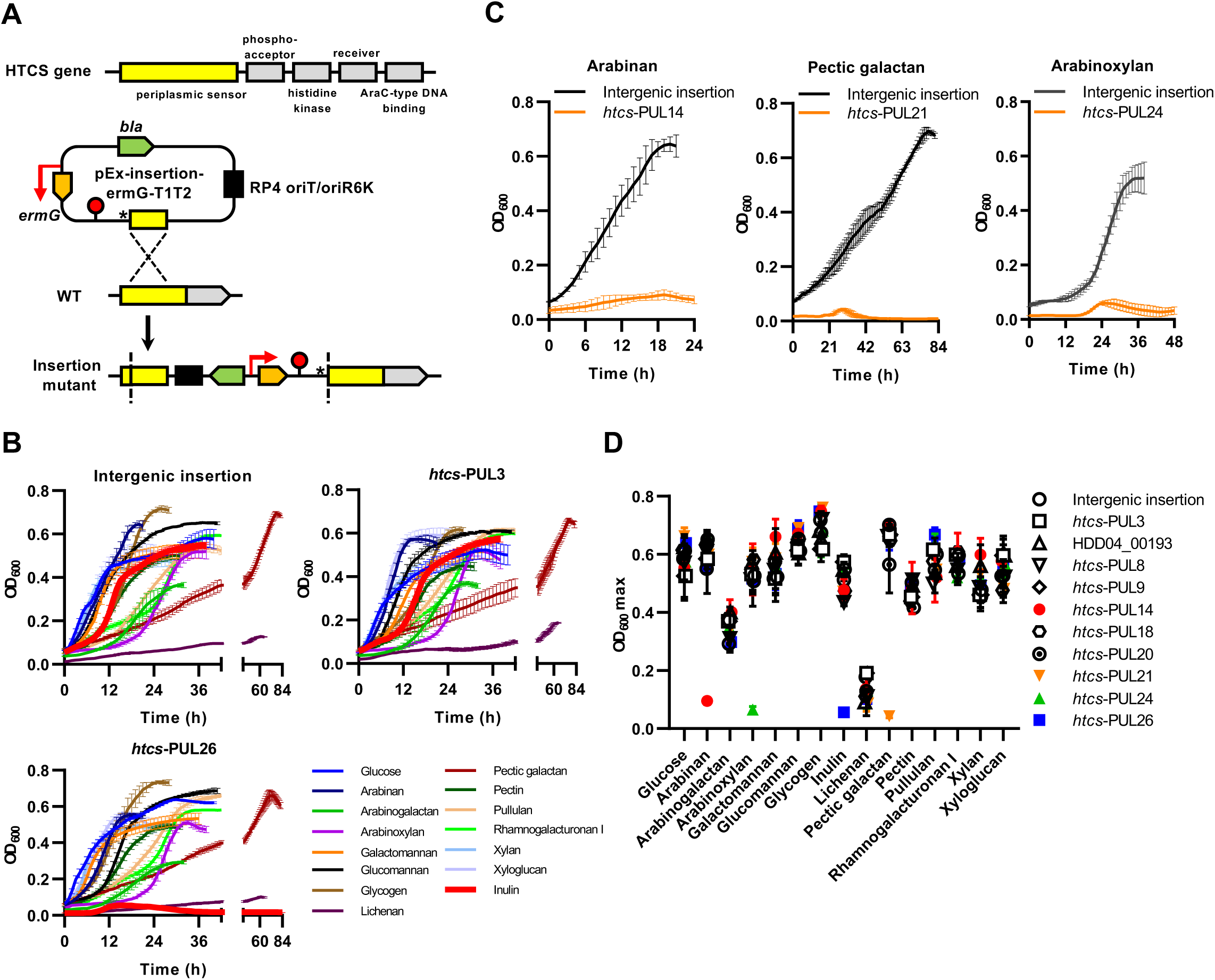
Identification of HTCS and associated PULs essential for utilization of polysaccharides using targeted gene inactivation. (A) Schematic illustration for gene inactivation strategy targeting HTCS gene candidates in *P. copri* HDD04. (B) Growth of *P. copri* HDD04 strains with plasmid insertions at an intergenic region (control) and two representative putative HTCS genes, respectively, in minimal media (MM) supplemented with glucose or indicated polysaccharides. (C) Growth of *P. copri* HDD04 strains with respective *htcs*-PUL14, -PUL24, and -PUL24 insertion compared to an intergenic insertion mutant on arabinan, pectic galactan, and arabinoxylan, respectively. (D) Maximum growth (OD_600_ max) of mutant strains (n=11) in MM supplemented with glucose or indicated polysaccharides. Error bars represent the standard error of the means (SEMs) in (B) and SDs in (C) and (D) of the biological replicates from three carbohydrate arrays with each carbohydrate tested in duplicate, respectively.

Together, these experiments not only demonstrate the utility of the gene inactivation strategies to perform functional studies in *P. copri*, but also uncovered the link between PUL associated regulatory genes and metabolic phenotypes on utilizing specific polysaccharides for *P. copri*.

### Construction of an allelic exchange system for validating the function of PUL regulators on polysaccharide degradation

Although our genetic insertion system is efficient in generating mutants for rapid determinations of phenotypes, it some limitations: (1) the selective pressure provided by antibiotics is constantly required in the medium for plasmid integrants with corresponding antibiotic resistance markers; (2) it would be challenging to target relatively smaller genes because the yield of conjugants is negatively correlated with the homology arm cloned in the conjugative plasmid as shown in Figure 1E; (3) the integration of the plasmid may cause a polar effect on the expression of downstream genes, especially complicating the characterization of each gene’s role in an operon. Therefore, we aimed to establish an allelic exchange system for unmarked gene deletion and complementation in *P. copri*. One of the most widely used system for allelic exchange in bacteria is based on the levansucrase gene (*sacB*), which catalyzes the hydrolysis of sucrose and synthesizes the toxic compound levan (Gay et al., 1985; Recorbet et al., 1993). Two key criteria have been identified to limit its application: (1) The inability of bacteria to grow properly on agar plates in the presence of relatively high concentrations of sucrose and (2) whether the expression of *sacB* can effectively select gene deletion mutants in the presence of sucrose.

Hence, we first determined the growth of *P. copri* DSM18205 and HDD04 on agar plates with increasing concentrations of sucrose. As media base, we employed yeast extract and tryptone (YT) supplemented with horse blood instead of BHI to reduce salt concentrations, which have been demonstrated to decrease the sucrose sensitivity of *E. coli* (Blomfield et al., 1991) and have been observed by us to interfere with *P. copri* growth on high sucrose concentration (data not shown). Both strains showed the normal colony numbers and morphology until a sucrose concentration of 6%, while *E. coli* tolerated up to 10% sucrose (Figure S3A). The inability to grow under these conditions was likely caused by osmotic pressure, as similar results were obtained for *P. copri* strains with increasing the concentration of glucose in YT+blood agar (Figure S3B). In order to ensure the selectivity of sucrose without affecting growth of *P. copri*, a working concentration of 5% sucrose was chosen. Notably, 5/10 strains were not able to grow in the YT+blood media in absence of sucrose reflecting their distinct nutritional requirements compared to other strains (Figure 3B and S3C).

**Figure 3.**
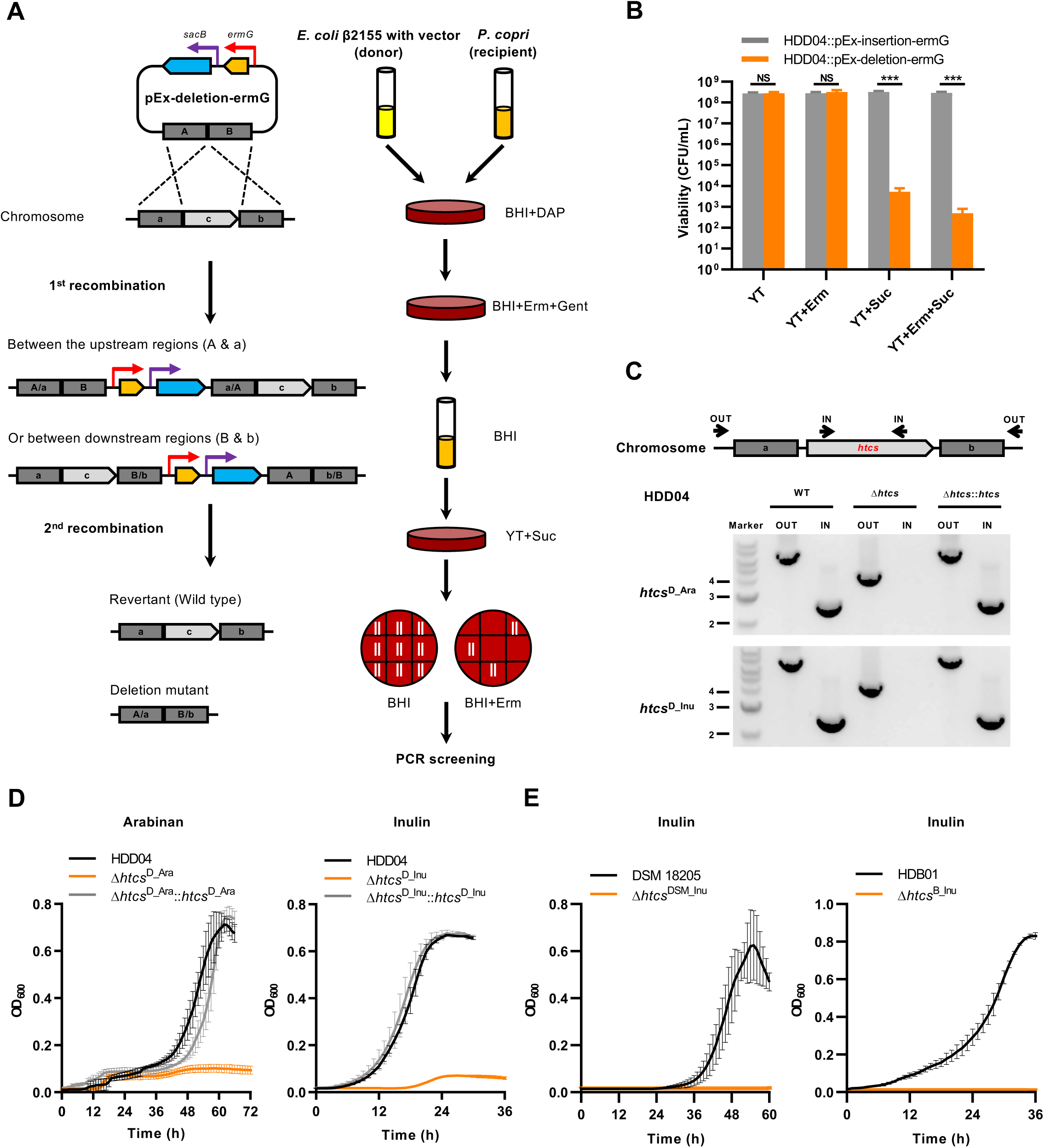
Development of a conjugation-based gene deletion and complementation platform for *P. copri* strains. (A) Schematic illustration for allelic exchange using pEx-deletion-ermG. (B) Viability of *P. copri* HDD04 with integration of pEx-insertion-ermG or pEx-deletion-ermG into the chromosome on YT agar plates with indicated supplements. Error bars represent the mean of three biological replicates ± SDs (***p < 0.001). (C and D) Generation of *P. copri* HDD04 Δ*htcs*^Ara^ and Δ*htcs*^Inu^ and quantification of growth. Detection of deletion and complementation of two HTCS genes in *P. copri* HDD04 by PCR using “OUT” and “IN” primer pairs (C) and growth of indicated strains in arabinan or inulin (D), respectively. (E) Growth of the wild-type and *htcs*^Inu^ mutant strains of DSM 18205 and HDB01, respectively. In (B, D, and E), the data represent the means of three biological replicates ± SDs.

Next, a derivative vector of pEx-insertion-ermG named pEx-deletion-ermG was created by joining the promoter of *gdhA* gene to a promoterless copy of *sacB* and inserting it downstream of *ermG* (Figure 3A and S2D). The homology arm for targeting the *bgl* gene (Figure 2A) was cloned into the pEx-deletion-ermG, resulting in pEx-deletion-ermG-HDD04-bgl. We individually integrated the pEx-deletion-ermG-bgl and pEx-insertion-ermG-bgl into HDD04 (Figure 3B). Erm^R^ colonies were readily obtained for both plasmids and displayed normal colony morphology as wild type, indicating that the expression of *sacB* did not affect the growth of *P. copri* in the absence of sucrose (Figure 3B). Plating these Erm^R^ colonies containing pEx-deletion-ermG-bgl in the presence of sucrose significantly reduced CFU by 10^4^-fold (Figure 3B). In contrast, the same strain carrying pEx-insertion-ermG-*bgl* exhibited equivalent viability in the presence and absence of sucrose. These results illustrated that expression of *sacB* effectively functioned as sucrose-based selection.

We subsequently assessed the false positive rate of this counterselection system by plating Erm^R^ on YT+Erm+Suc plates and found that approximately 1 out of 6.7×10^5^ cells in the bacterial population were Suc^R^ but still Erm^R^, i.e. carried the plasmid. Of note, 50 from 500 colonies were randomly picked and restreaked on YT+Erm and YT+Suc plates to evaluate whether the Suc^R^ phenotype of these “escapers” was due to genetic mutations. Unexpectedly, they all showed Erm^R^ but Suc^S^ phenotypes. We further sequenced the *sacB* gene and its promoter sequences in 10 random-selected clones, yet none of them had mutations. This suggested that the Suc^R^ phenotype of these escapers were attributable to phenotypic but not genetic causes. Similar level of selectivity in the *sacB*-sucrose based system was recapitulated in other five *P. copri* strains (Figure S2D, S2E, and S3D). Taken together, these results demonstrate the utility of the *sacB*-based counterselection system for targeted allelic exchange in *P. copri*.

### Genetic deletion and complementation demonstrate essential function of HTCS genes on degrading plant polysaccharides

As a proof of concept, we chose two HTCS genes, *htcs*^D_Ara^ and *htcs*^D_Inu^ as our targets for genetic deletion and complementation. A schematic for describing the allelic exchange methodology during gene-editing process including plasmid construction, allelic exchange, and mutant selection is presented in Figure 3A. In brief, we (1) cloned up- and down-stream regions of the target gene into the pEx-deletion-ermG plasmid and transferred the plasmid into *E. coli* β2155; (2) performed conjugation between *P. copri* and *E. coli* carrying the plasmids followed by selection of plasmid integrants (1^st^ recombination); (3) passaged the Erm^R^ clones without any selection, permitting the spontaneous allelic exchange (2^nd^ recombination); (4) carried out the selection of bacteria that had lost the plasmid (revertant and deletion mutant); (5) validated the Erm^S^ phenotype of selected clones and screened for the clones with gene deletion mutations. Of note, PCR screening and Sanger sequencing of Erm^S^ clones obtained after the counterselection step revealed that 37.5% to 56.3% of clones are confirmed deletion mutants (Table S4) with all remaining clones being revertants, showing the precise performance of our targeting system. Following these procedures, *htcs*^D_Ara^ and *htcs*^D_Inu^ were successfully deleted in HDD04, generating Δ*htcs*^D_Ara^, and Δ*htcs*^D_Inu^ gene deletion strains accordingly (Figure 3C). We subsequently complemented the deletion mutants with the corresponding HTCS genes, respectively, through a similar genetic procedure except that we cloned the target gene and its flanking regions into pEx-deletion-ermG (Figure 3C). In line with our previous findings in Figure 2D, the HTCS-deficient mutants failed to grow on their previously identified substrates, while complementation of HTCS genes in the mutant strains restored the growth to the levels of the wild type HDD04 strain (Figure 3D).

To demonstrate that the allelic exchange system can be also applied to *P. copri* strains with relatively lower yields of transconjugants, individual deletion of homologous *htcs*^D_Inu^ genes were performed in the DSM 18205 (*htcs*^DSM_Inu^, DSM18205_02724) and HDB01 (*htcs*^B_Inu^, HDB01_02906) strains (Figure S3E). Similarly, deletion of *htcs*^D_Inu^ displayed dramatic grow defects in both DSM 18205 and HDB01, respectively (Figure 3E).

In conclusion, these results demonstrate the utility of our novel allelic exchange system in the type strain and other *P. copri* isolates for establishing a causal relationship between genotypes and phenotypes, as best exemplified by specific HTCS genes and the growth phenotypes on arabinan and inulin.

### Plant-derived polysaccharides induced the transcription of distinct PUL-associated genes in *vitro* and in *vivo*

To determine whether polysaccharides induce the expression of specific PULs associated with the identified HTCS genes or rather broader changes in multiple PULs, we performed transcriptome profiling of *P. copri* HDD04 cultures grown in MM supplemented with either glucose or one of four plant polysaccharides as the sole carbohydrate (Table S5). As we expected, the *susC/D*-like elements in PUL14, 21, 24, and 26 exhibited the largest upregulation upon their respective polysaccharide substrates compared to MM+Glucose (Figure 4A), further confirming our genetic characterization (Figure 2 and 3). We therefore defined these four PULs as PUL^D_Ara^, PUL^D_PecGal^, PUL^D_AraXyl^, and PUL^D_Inu^ (Figure 4B). Using average fold-change of *susC/D*-like genes in each PUL as reference, PUL^D_Inu^ showed a relatively low (3.3-fold) upregulation in response to inulin, whereas PUL^D_Ara^, PUL^D_PecGal^, and PUL^D_AraXyl^ displayed much higher induction levels (PUL^D_Ara^: 266.1-fold; PUL^D_PecGal^: 983.7-fold; PUL^D_AraXyl^: 159.3-fold; Figure 4A). It is worth noting that the disruption of HTCS gene in PUL24 resulted in the growth deficiency on arabinoxylan, but did not affect the growth pattern on xylan (Figure 2C and Figure 4A). This is in disagreement with a previous prediction, which was based on bioinformatic analysis and growth assays, of a PUL24 homolog being a xylan processing PUL (Figure S4A) (Fehlner-Peach et al., 2019). Interestingly, genome-wide only *susC/D*-like genes from PUL^D_Inu^ were significantly upregulated, indicating an extremely specific system for processing inulin by *P. copri*. Yet, multiple *susC/D*-like elements were induced (>10-fold change) by the other three plant-derived polysaccharides (Figure 4A). For instance, *susC/D*-like pairs in PUL^D_PecGal^ (PUL24, 23.9-fold) were also greatly expressed when cells were exposed to arabinan. The transcriptional response of *susC* homologs in four identified PULs to each tested polysaccharide were further validated by real-time quantitative PCR (RT-qPCR) (Figure 4C). Hence, we hypothesized that the multiple PULs are response to the degradation products of plant polysaccharides, e.g., PUL^D_PecGal^ was likely induced by the degraded components of pectin fragment attached to the main-chain arabinofuranosyl residues of sugar beet arabinan (Figure S4B). Surprisingly, PUL15 (HDD04_02377-87) possesses genes encoding GH13 and GH97, which were previously demonstrated to degrade various glucans (Cerqueira et al., 2020; Koropatkin et al., 2008), was commonly upregulated by arabinan, pectic galactan, and arabinoxylan. Besides the regulation of PUL-associated genes, there were also genes encoding polysaccharide catabolism enzymes that displayed >10-fold upregulation by the polysaccharides, e.g. one putative extracellular exo-alpha-L-arabinofuranosidase precursor gene in MM+Arabinan (HDD004_02362, Table S5). Similarly, a putative gene operon was strongly upregulated in response to arabinan and arabinoxylan, which has a high similarity to the arabinose utilization system in *B. thetaiotamicron* (Schwalm et al., 2016) (Figure S4D).

**Figure 4.**
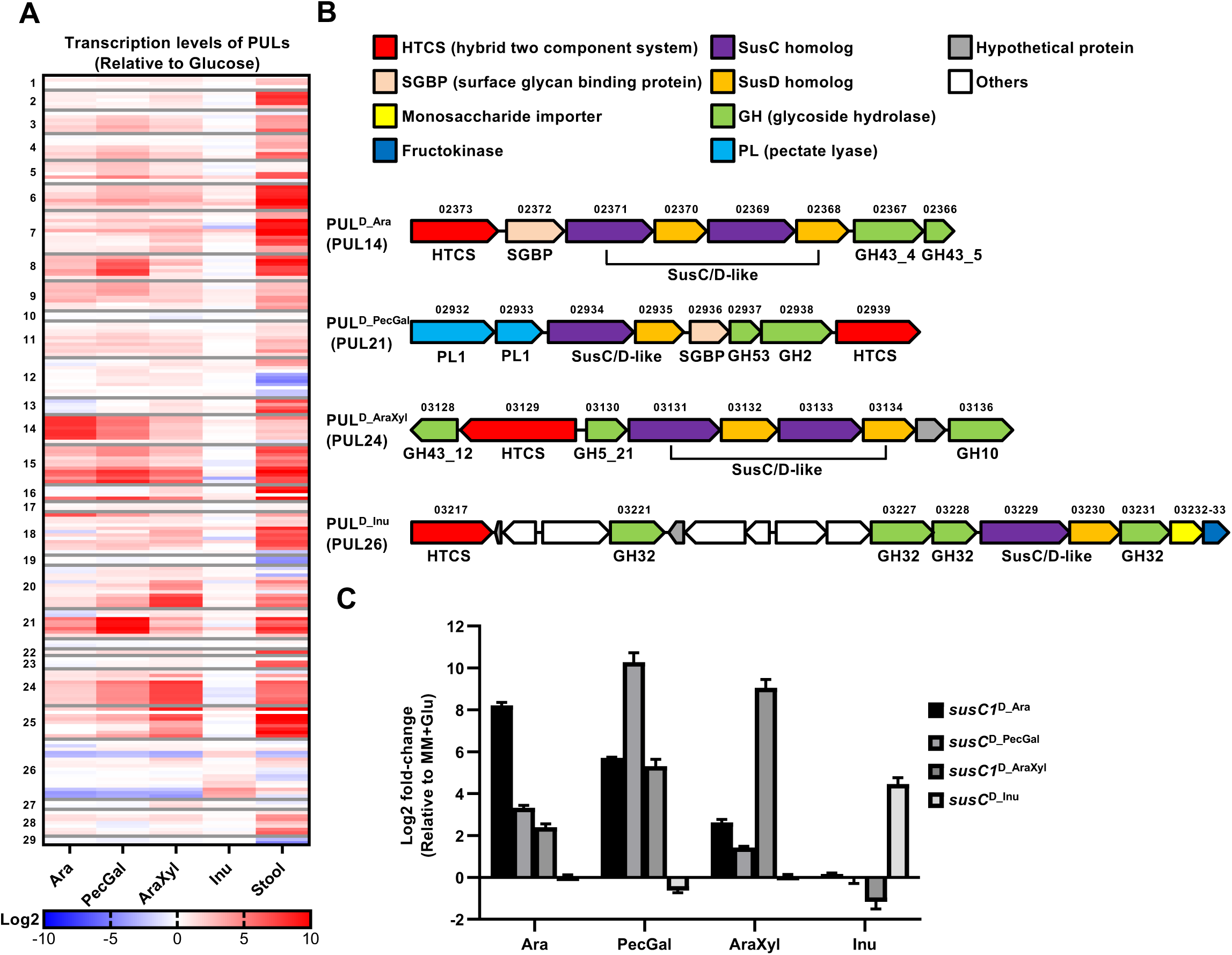
Transcriptional adaptation of *P. copri* HDD04 to distinct plant polysaccharides and in the human gut. (A) Heatmap showing the induction of *susC* and *susD* homologs in each predicted PUL from *P. copri* HDD04 in MM supplemented with indicated polysaccharides or in a fecal sample. Average gene expression of every *susC/D* element (two or four genes) was calculated and normalized to expression in glucose. The heatmap shows the average log2 fold change of *susC/D* pairs within the predicted PULs. PUL14 (PUL^D_Ara^), PUL21 (PUL^D_PecGal^), PUL24 (PUL^D_AraXyl^), and PUL26 (PUL^D_Inu^) are highlighted by black borders. (B) Genetic architectures of PUL^D_Ara^, PUL^D_PecGal^, PUL ^D_AraXyl^, and PUL^D_Inu^, in *P. copri* HDD04. Genes in PULs are annotated by their gene numbers and predicted functions.. (C) *In vitro* transcriptional response of targeted *susC* homologs in MM+indicated polysaccharide in comparison with MM+Glucose reference.

To identify whether *P. copri* actively utilizes these PULs *in vivo*, a metatranscriptome analysis was performed from a stool sample collected from the donor, from which HDD04 was isolated. Strikingly, except PUL^D_Inu^ that displayed a slightly increased transcription compared to MM+Glucose (average of fold-change *susC/D*-like genes: 1.2-fold), the *susC/D* homologs in the other three identified PULs (PUL^D_Ara^: 4.2-fold; PUL^D_PecGal^: 215-fold; PUL^D_AraXyl^: 44.6-fold) were actively expressed. Moreover, 16 other PULs displayed upregulation from 2.36-fold (PUL1) to more than 8000-fold change (PUL25), which suggests additional substrates from the human diet can be targeted by various PULs in *P. copri* (Figure 4A). Collectively, these analyses illustrate that *P. copri* carries out an efficient and diverse polysaccharide processing by orchestrating its associated gene expression profile *in vitro* and *in vivo*. Further functional gene studies will be required to understand which polysaccharides can be utilized *in vivo* by *P. copri*.

### PUL^D_Ara^, PUL^D_PecGal^, PUL^D_AraXyl^ and PUL^D_Inu^ are conserved among genetically diverse *P. copri* strains

Recent studies reported that *P. copri* isolates exhibited extensive genomic and phenotypic variations (Fehlner-Peach et al., 2019; De Filippis et al., 2019; Tett et al., 2019). To examine whether utilization of arabinan, pectic galactan, arabinoxylan, and inulin is a common capacity based on the genetic content of *P. copri* species, we performed a comparative genomic analysis to identify corresponding PULs in strains from our *P. copri* strain collection. Notably, PULs carrying homologous HTCS/SusC genes compared to HDD04 were found in each of the *P. copri* strains (Table 1 and S3; Figure S5). The gene organization and content of these PULs varied from conserved, e.g. PUL^AraXyl^, to variable, e.g. PUL^Inu^ (Figure S5). Hence, we next determined the growth in MM supplemented with arabinan, pectic galactan, arabinoxylan, or inulin. Most *P. copri* strains grew on the tested polysaccharides with the exception of HDA03 that could not grow on pectic galactan and arabinoxylan (Figure S5). Strikingly, genetic evidence potentially explaining the inability to use specific polysaccharides could be easily identified (Figure S5). Specifically, natural mutations in the HTCS genes of PUL^AraXyl^, i.e. a truncation of HTCS^AraXyl^, and of PUL^PecGal^, i.e. deletion, are responsible for the “no growth” phenotypes of HDA03 on these polysaccharides. Additionally, the cognate first *susC* gene of PUL^AraXyl^ as well as the SGBP gene of PUL^PecGal^ were truncated into two segments in HDA03, which could further contribute to the inability to utilize these polysaccharides.

**Table 1.** Gene organizations of two types of arabinan processing PULs in multiple *P. copri* and two *Bacteroides* type strains with the capacities of growing on arabinan.

| Clade | Strain | OD<sub>600</sub> max | PUL organization | Type |
| --- | --- | --- | --- | --- |
| A | DSM 18205 | 0.446±0.066 |  | I |
| A | HDA03 | 0.720±0.028 |  |  |
| A | HDA04 | 0.639±0.026 |  |  |
| A | HDC01 | 0.505±0.058 |  |  |
| A | RHA03 | 0.597±0.039 |  |  |
| A | RPA01 | 0.552±0.071 |  |  |
| C | HDE04 | 0.788±0.008 |  |  |
| C | HDE06 | 0.906±0.035 |  |  |
| D | HDD05 | 0.676±0.019 |  |  |
| / | *P. vulgatus* ATCC 8482 | / |  |  |
| A | HDD04 | 0.731±0.028 |  | II |
| A | HDB01 | 0.587±0.044 |  |  |
| E | HDD12 | 0.731±0.052 |  |  |
| / | *B. theta* VPI-5482 | / |  |  |

Notably, the PUL^PecGal^ in HDE04 and HDD12 and PUL^Inu^ in HDD05 and HDD12 do not contain *susC/D*-like elements, suggesting that they are non-essential for utilizing pectic galactan and inulin in these host strains. These cases of “incomplete” PULs highlight the limitations of using *susC/D* homologs as genetic markers to predict the growth phenotypes on specific polysaccharides substrates.

### Two types of arabinan processing PULs in *P. copri* display clade-specific distribution and diet-dependent expansion in the human gut microbiome

While functionally all tested strains were able to use arabinan, we found that the arabinan processing PULs genetically displayed two distinct structures among the 12 strains (Table 1). Specifically, two strains from clade A and the single strain from clade E feature an almost identical gene organization with a SGBP-like gene in front of two pairs of *susC/D*-like genes, whereas the remaining strains from clade A, as well as the strains from clade C and D encode a single pair of *susC/D*-like genes followed by a SGBP-like gene. In the following we refer to two types of PULs as type-I (single copy) and type-II (tandem repeat), and the two *susC/D*-like pairs in type-II PUL^Ara^ as *susC1*, *susD1*, *susC2*, and *susD2*, respectively. Of note, similar PUL^Ara^ types have been noticed in *Phocaeicola vulgatus* (formerly *Bacteroides vulgatus*) with type-I and *B. thetaiotaomicron* with type-II (Table 1) (Lynch and Sonnenburg, 2012; Martens et al., 2011; Patnode et al., 2019).

Phylogenetic analysis of protein sequences encoded by HTCS genes from the 12 *P. copri* strains as well as *P. vulgatus* ATCC 8482 and *B. thetaiotaomicron* VPI-5482 shows a clade-driven evolutionary pattern, which closely resembled that of genome-based tree of the *P. copri* complex (Figure 1A). The inability of the mutant strain DSM18205 *htcs*^DSM_Ara^ to grow on arabinan validates that the HTCSs of two arabinan utilizing systems are functionally conserved in the two types of PUL^Ara^ (Figure 5B). In contrast, the *P. copri*-derived SusC and SusD homologs form three distinct evolutionary branches in the tree that differ from the proteins in *B. thetaiotaomicron* and *P. vulgatus,* but still show a relative similarity of these proteins corresponding to the PUL^Ara^ types, respectively (Figure 5A and S6A). Similar to the *susC/D*-like genes, the SGBP-like proteins are clustered by PUL^Ara^ type rather than *P. copri* clade (Figure S6B). We next performed functional studies complementing the phylogenetic analysis. Because the *susC1* gene but not the *susC2* gene is required in *B. thetaiotaomicron* for growth on arabinan as described previously (Luis et al., 2018), the genes encoding *susC* in type-I, and *susC1* and *susC2* in type-II system of *P. copri* were individually in-frame deleted to explore their necessity in *P. copri* (Figure 5B). In agreement with previous observations in *B. thetaiotaomicron*, only *susC1* but not *susC2* is essential for type-II PUL^Ara^ carrier HDD04 (Figure 5B). Moreover, deletion of *susC* in type-I PUL^Ara^ abolished the growth capacity of *P. copri* DSM 18205 on arabinan (Figure 5B). These results indicated that *P. copri* strains encode highly similar sensory/regulatory systems for sensing arabinan-derived ligands and transcriptional activation of PUL^Ara^, but that PUL^Ara^ encodes distinct modules, i.e. *susC*-*susD*-*SGBP* in type-I and *SGBP*-*susC1*-*susD1-susC2*-*susD2* in type-II, for carbohydrate binding and importing.

**Figure 5.**
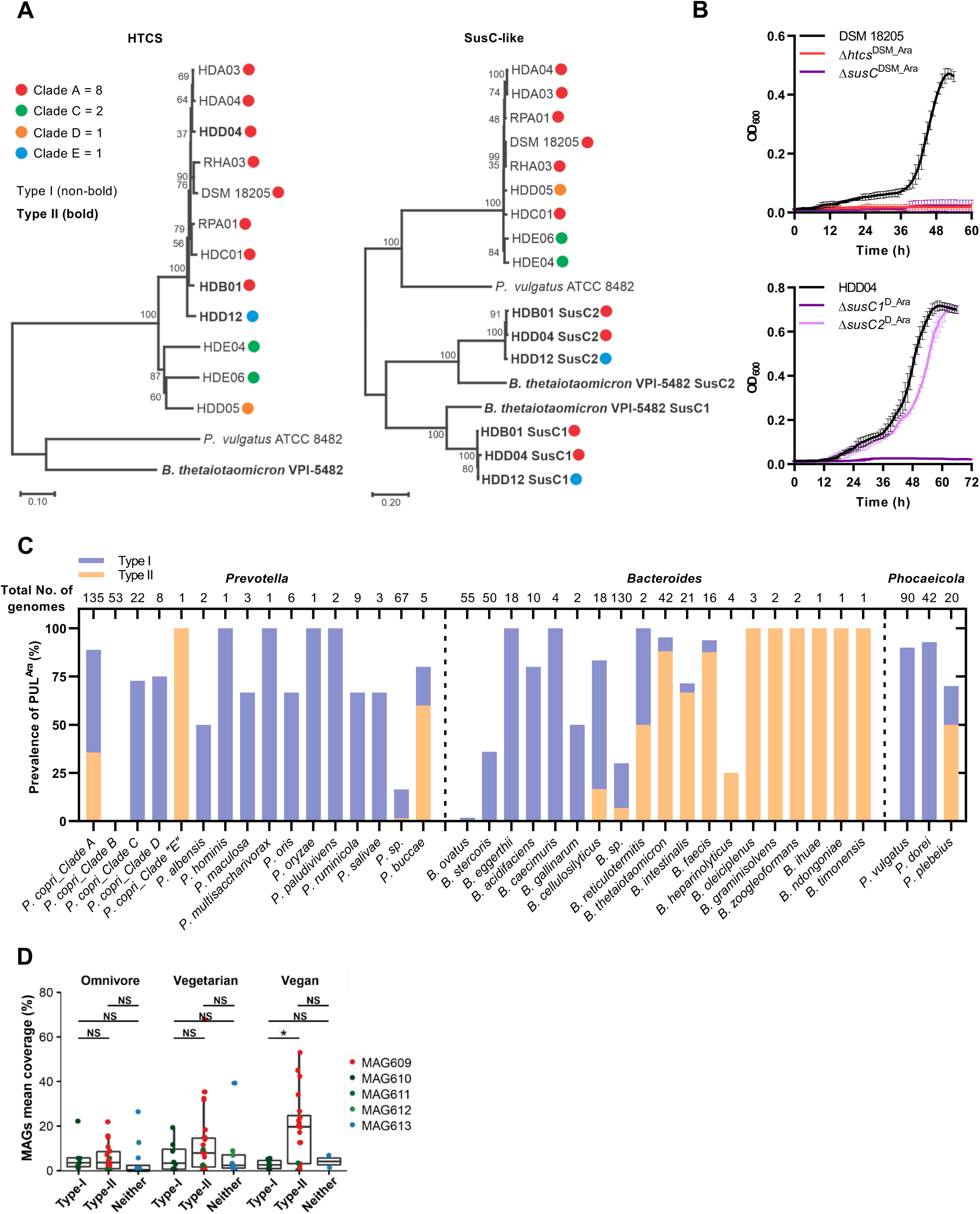
Genetic and phylogenetic analyses of PUL^Ara^ types in members of the genera *Prevotella, Phocaicola* and *Bacteroides*. (A) Phylogenetic trees of HTCS and SusC-like proteins encoded by two types of arabinan processing PULs in diverse *P. copri* strains, *B. thetaiotaomicron* VPI-5482, and *P. vulgatus* ATCC 8482. The isolates from distinct *P. copri* clades are indicated by dots in different colors. The proteins from type-II PUL^Ara^ are highlighted in bold. (B) Growth of *P. copri* DSM 18205 and HDD04 wild-type strains and indicated mutants in MM+Arabinan. (C) Distribution of two types of arabinan processing PULs in the members of genera *Prevotella*, *Bacteroides* and *Phocaeicola*. Total number of genomes for each clade or species group that were analyzed is indicated above the bars. (D) The association between the two types of PUL^Ara^ in *P. copri* and host dietary preference. The relative abundance of identified *P. copri* MAGs for each individual was grouped based on the presence or absence of two types of PUL^Ara^ in each dietary habit. Asterisks indicate Wilcoxon U-test significant differences (*p < 0.05; NS, p > 0.05, not statically significant)

To gain a broader understanding of the prevalence of PUL^Ara^ types in *P. copri* as well as related *Prevotella* spp., *Bacteroides* spp. and *Phocaeicola* spp., PUL^Ara^ were predicted from 1602 non-redundant genomes retrieved from the NCBI genome database (n=1504) and from a recent comprehensive metagenomic *P. copri* survey (n=98) (Tett et al., 2019). Together with our strains (n=12), we identified that 499 out of 1614 genomes encode either type-I or type-II PUL^Ara^, suggesting that the arabinan utilization potential is frequently found in these genera but not ubiquitous (Figure 5C and Table S6). In agreement with the results from the analysis of our limited strain collection, type-I PUL^Ara^ is present in *P. copri* clades A (55.3%), C (72.7%) and D (75%), while the type-II is encoded by *P. copri* strains from the clade A (36.8%) and the single strain from the clade E (HDD12) (Figure 5C). Notably, none of the two types as well as the arabinose utilization operon we identified was found in any of the 53 screened genomes of clade B strains (Figure 5C), which is consistent with previous reports that this clade lacks the genes and capacity for arabinose and arabinan utilization (Tett et al., 2019). The two types of PUL^Ara^ are also widespread in other members of the *Prevotella* spp., *Bacteroides* spp. and *Phocaeicola* spp. (Figure 5C). Notably, our extended analysis shows that while some species displayed either a type I- or type II-dominated distribution, e.g., *P. vulgatus* (n=90 genomes, type-I: 90%, type-II: 0%), other species such as *Phocaeicola plebeius* (n=20 genomes, type-I: 20%, type-II: 50%) and *B. thetaiotaomicron* (n=42 genomes, type-I: 7.14%, type-II: 88.1%) displayed both types of arabinan utilization systems similar to *P. copri* species (Figure 5C). As the evolutionary analysis of *Bacteroides* spp. and *Phocaeicola* spp. is limited, these dominations could also result from the clade-specific distribution as *P. copri*. Interestingly, only 1 out of 55 *Bacteroides ovatus* genomes carry type-I PUL^Ara^, which not only suggests the occurrence of horizontal gene transfer of type-I PUL^Ara^ between *B. ovatus* and other type-I PUL^Ara^ carriers (Figure 5C), but also provides an explanation for the apparent lack of arabinan utilization in most *B. ovatus* strains (Martens et al., 2011).

The increased relative abundance of *P. copri* in the gut microbiota has been associated with fiber-rich diets (De Filippo et al., 2010; Fragiadakis et al., 2019; Ruengsomwong et al., 2016; Wu et al., 2011). As the *in vitro* results described above suggest a potential contribution of the PUL^Ara^ for fitness advantage within the ecosystem, we investigate whether *P. copri* encoding different types of PUL^Ara^ displays a diet-modulated abundance in the human gut. Therefore, we performed a specialized analysis of a publicly available dataset from one recent study comparing the differences in the gut microbiota of individuals consuming omnivore, vegetarian, and vegan diet (De Filippis et al., 2019). In a previous study, we identified five distinct *P. copri* metagenome-assembled genomes (MAGs) from four clades in individuals of that cohort (Figure 5D). MAG610 (clade C) encodes a type-I PUL^Ara^, whereas two MAGs 609 and 611 (clade A and C) carry the type-II counterpart. Two MAGs 612 and 613 (clade C and B) contain neither of the PULs (Figure 5D). The relative abundance of each MAG was then calculated per individual and grouped based on the presence of type-I or type-II PUL^Ara^ in the individuals with three distinct dietary preferences (Figure 5D). While there was no significant difference based on PUL^Ara^ presence and type in omnivore and vegetarian diets, remarkably, in vegans, PUL^Ara^ type-II positive MAGs showed higher abundance than those with type-I PUL^Ara^ (Figure 5D). This suggests that *P. copri* strains specifically benefit from type-II PUL^Ara^ under particular dietary conditions. Further genetic and functional characterizations will be required to further understand the differential precise nature of two PUL^Ara^ systems. Yet, taken together our comprehensive analyses illustrate the importance of arabian utilization systems for *P. copri* fitness *in vitro* and *in vivo*.

## Discussion

Studying the biology of many human commensals is hindered by diverse obstacles, e.g. challenging cultivation, extensive strain-level variation, low number of publicly available strains, and the lack of genetic tools (Fehlner-Peach et al., 2019; De Filippis et al., 2019; Ley, 2016; Tett et al., 2019). In this study, we developed a genetic toolkit that allows versatile genetic manipulations for a wide range of *P. copri* strains, and applied our genetic platform coupled with genomic, transcriptomic, and phenotypic approaches to provide insights into the genetic basis of polysaccharide utilization of this prevalent gut bacterium. Particularly, the recognition that *Prevotella* spp. including *P. copri* are a dominant part of the “non-westernized” microbiome as well as their unexplained antagonism with *Bacteroides* spp. has elevated the interest in the members of this genus (Arumugam et al., 2011; Costea et al., 2017; Johnson et al., 2017; Ley, 2016). While a series of genetic tools have been created for the genus *Bacteroides* (Bencivenga-Barry et al., 2020; García-Bayona and Comstock, 2019; Goodman et al., 2011; Koropatkin et al., 2008; Lim et al., 2017; Mimee et al., 2015), none have been reported for *P. copri* despite its diverse associations to human diseases. Moreover, few functional genetic studies reported on targeting genes in animal-derived *Prevotella*, i.e. *Prevotella ruminicola* (Gardner et al., 1996; Ogata et al., 1999; Shoemaker et al., 1991) and *Prevotella bryantii* (Accetto and Avguštin, 2007; Accetto et al., 2005). Together, this suggests the existence of limitations preventing the simple transfer of established genetic tools to *P. copri*.

Here, by redesigning and optimizing the key genetic elements in the conjugative plasmids and experimental procedures, we developed an anaerobic conjugation based system, which overcomes the genetic intractability of diverse *P. copri* strains. For instance, the optimization of promoter strength for the selection marker resulted in a 306.6-fold increase in conjugation outcomes. Another key factor was the evaluation of different donor and recipient strains improving conjugation outcome by 83.7-fold and up to 14,160-fold, respectively. Of note, naturally occurring antibiotic resistances can complicate gene targeting, but the combination of plasmids carrying either *ermG* or *tetQ* antibiotic markers allowed the generation of insertion mutants for 11 out of 12 *P. copri* strains in our strain collection representing four distinct clades.

In a proof-of-concept of our approach, we focused on one strain, i.e., HDD04, with high conjugation capacity and robust growth in chemically defined medium, targeting ten HTCS regulators for controlling PUL expression, and identified four HTCS genes with essential functions in utilizing plant polysaccharides. Next, we adapted the *sacB*-sucrose system to *P. copri* that enabled the selection of mutants after allelic exchange with a 3.1×10^−7^ to 4.9×10^−6^ false positive rate. This efficacy is similar to the that obtained for a recent system used for allelic exchange in *Bacteroides* spp. (Bencivenga-Barry et al., 2020). Of note, compared to other counterselection systems, such as the genes encoding 30S ribosomal protein S12 (*rpsL*) for Proteobacteria and thymidine kinase (*tdk*) for *Bacteroides* spp. (Dean, 1981; Koropatkin et al., 2008; Reyrat et al., 1998), the *sacB*-sucrose system does not require any genetic modification of the recipient strain in advance. The utility of the *sacB*-sucrose system was demonstrated by deletion of HTCS genes in three strains, including ones with relatively lower conjugation capacity. Despite these advantages, the allelic exchange system for *P. copri* does have limitations, e.g. it requires multiple selection steps and the ability to tolerate the osmotic pressure of high sucrose concentrations. Other counterselection markers, such as inducible antibacterial effectors (Bte1 and Bfe1) utilized in *Bacteroides* species (Bencivenga-Barry et al., 2020; García-Bayona and Comstock, 2019), can be envisioned, yet it has been noticed that *P. copri* DSM 18205 was not affected by the Bfe1 effector (Chatzidaki-Livanis et al., 2016). Another limitation relates to complementation of gene deletions, e.g. in *B. thetaiotaomicron*, genetic complementation is accomplished by integrating a plasmid carrying the complemented gene into the chromosome (Koropatkin et al., 2008; Wang et al., 2000). Specifically, the integration vector pNBU2 encodes a tyrosine integrase, which mediates sequence-specific recombination between the attN site of pNBU2 and one of two attBT sites located in the 3’ ends of the two tRNASer genes on the *B. thetaiotaomicron* chromosome. Because no identical or similar attachment DNA sequences were found in *P. copri*, we so far carried out the complementation by the same allelic exchange approach as used for gene deletion. Thus, development of a similar integration vector for *P. copri* or a *P. copri* parent strain carrying attachment sites for pNBU2, has the potential to simplify the process of genetic complementation.

The high prevalence and increased abundance of *P. copri* in the intestinal microbiota is frequently associated with consumption of fiber-rich diets, which has inspired the research for underlying genetic basis of polysaccharide utilization in *P. copri*. Combinations of comparative genomics and phenotypic assays have predicted the substrates for the PULs harboring well-defined CAZymes-coding genes (Fehlner-Peach et al., 2019). However, the direct contribution of specific PULs for polysaccharide substrates has not been documented for *P. copri*. Our genetic studies confirmed the previous bioinformatic prediction of arabinan as the substrate for the PUL14 homologs (Fehlner-Peach et al., 2019), Additionally, three new PUL/polysaccharide combinations were identified in *P. copri*. It is intriguing that PUL24, whose homologous gene cluster was predicted based on bioinformatic analysis as xylan-processing PUL, was identified in this study to be essential for wheat arabinoxylan utilization, but non-essential for xylan utilization. Thus, genetic approaches coupled with phenotypic and transcriptome analyses present a framework for a more accurate characterization of PULs. Notably, the gene organization and content of PULs varied between *P. copri* strains from conserved to variable. For instance, the strictly conserved synteny of the arabinan and arabinoxylan processing PULs suggests that they have been under the positive selection pressure, as discussed below. In contrast, the gene content of the PULs for pectic galactan and inulin are relatively variable, suggesting the non-essentiality of certain genes for supporting growth of *P. copri* on these two carbon sources, including even *susC/D*-like genes. These observations support the model that *susC/D*-like element in some but not all PULs are essential for the uptake of polysaccharides and are therefore suboptimal markers for carbohydrate utilization potential. Finally, *P. copri* strain carry either one of two types of PUL^Ara^, i.e. type-I with a single *susC/D* pair or type-II PUL^Ara^ with tandem repeat *susC/D*, which appears to be shared phenomenon in the *Bacteroides*, *Phocaeicola* and *Prevotella* genera. In *P. copri*, the distribution of the two distinct PUL types showed largely clade-specific features, i.e. clade A encoded both types of PUL^Ara^, clades C and D only encoded type-I, and none of identified PUL^Ara^ was found in clade B (Fehlner-Peach et al., 2019; Tett et al., 2019). While the HTCS regulators independent of the PUL^Ara^ type shared high homology between members of the same clade, the *susC/D*-like and SGBP-like genes clustered by PUL type. Notably, PUL^Ara^ type-specific domination was observed in individuals consuming a vegan diet, suggesting the advantage of type-II over type-I in utilizing arabinan or potential other arabinose-based polysaccharides from dietary fibers in the human gut.

In summary, we have demonstrated the versatile capacities of the genetic toolbox for, firstly, generating a series of individual gene insertion mutants for phenotypic screening in parallel; secondly, enabling targeted gene deletion and complementation to establish causal relationship between genotypes and phenotypes; thirdly determining the impact of homologous genes in distinct *P. copri* strains on specific polysaccharide utilization. The toolbox will enable the dissection of more sophisticated biological interactions of *P. copri* with the human hosts during health and disease, such as investigate associations of *P. copri* to host metabolism in *vivo* (Kovatcheva-Datchary et al., 2015; Pedersen et al., 2016; De Vadder et al., 2016). Importantly, the platform was designed using general principles highlighting key technical details that can be modified and applied to other *Prevotella* species and even prominent bacterial genera from humans and other habitats. Moreover, these principles can be utilized for further development of high throughput genetic screening, such as transposon mutagenesis (Goodman et al., 2011) and CRISPRi (Peters et al., 2016), thereby advancing studies into systematically understanding the ecological and metabolic processes of microbiota and their impacts on host health and disease.

## STAR METHODS

### RESOURCE AVAILABILITY

#### Lead Contact

Further information and requests for resources and reagents should be directed to and will be fulfilled by the Lead Contact, Till Strowig.

#### Materials Availability

### KEY RESOURCES TABLE

#### Preparation of culture media for *Prevotella copri*

##### BHI+S liquid medium

The fetal bovine serum was heated at 56°C for 30 min to inactivate complement. 9.25 g of brain heart infusion (BHI) powder was dissolved in 225 mL double-distilled water (ddH_2_O) in 500-mL glass bottle. The medium was supplemented with 10% fetal bovine serum (FBS), and placed on the hotplate stirrer with 250°C for 20 min. The heated medium was then cooled down to room temperature, supplemented with 1 µg/mL vitamin K3, and filter-sterilized using a filter unit (0.22 mm pore diameter).

##### Minimal medium

Minimal medium (MM) was prepared as previously described with some modifications (Martens et al., 2008). It contained 100 mM KH_2_PO_4_ (pH 7.2), 15 mM NaCl, 8.5 mM (NH4)_2_SO_4_, 10 mL/L amino acid mix solution (250 mg each of L-alanine, L-arginine, L-asparagine, L-aspartic acid, L-cysteine, L-glutamic acid, L-glutamine, glycine, L-histidine, L-isoleucine, L-leucine, L-methionine, L-phenylalanine, L-proline, L-serine, L-threonine, L-tryptophan, L-tyrosine, L-valine, and 312 mg of L-lysine monohydrochloride into 1 L ddH_2_O), 10 mL/L purine/pyrimidine solution (200 mg each of adenine, guanine, thymine, cytosine, and uracil into 1 L ddH_2_O, pH 7.0), 10 mL/L ATCC Vitamin Mix, 10 mL/L ATCC Trace Mineral Mix, 100 μM MgCl_2_, 1.4 μM FeSO_4_·7H_2_O, 50 μM CaCl_2_, 1.9 μM hematin, 1 mg/L vitamin K3, 5 μg/L vitamin B12, 1 μL/L vitamin K1 solution, and 0.5 g/L cysteine. The commercial carbohydrates were prepared as described previously (Martens et al., 2011). Briefly, 10 g/L carbohydrate stock solutions (2× concentration) were sterilized by autoclaving at 121°C for 15 mins. When needed, 10 g/L carbohydrate solutions were added into 2× MM at a volume ratio of 1: 1.

##### YT+S agar

Sucrose was dissolved in ddH_2_O at 0.5 g/mL (50%) as a stock solution. 5 g yeast extract and 10 g tryptone were dissolved in 900 mL ddH_2_O. The resulting medium (YT) was autoclaved, cooled down to 50°C, and supplemented with 5% sucrose (100 mL 50% sucrose stock solution) for counter-selection for gene deletion and complementation of *Prevotella copri* strains. When necessary, the different volumes of sucrose solution were added in media.

##### Bacterial culture conditions

All strains, plasmids, and primers used are listed in Table S1. *Escherichia coli* strains were grown aerobically at 37°C on Luria-Bertani (LB) media. *E. coli* β2155 were specially cultured on LB supplemented with 0.3 mM 2,6-Diaminopimelic acid (DAP) (Demarre et al., 2005). *P. copri* was cultured in BHI+S liquid media, minimal media plus a carbon source, on BHI agar supplemented with 5% defibrinated horse blood or on YT agar supplemented with 5% defibrinated horse blood and 5% sucrose unless otherwise specified. Cultures were routinely grown and manipulated in an anaerobic chamber (Coy Laboratory Products) with an atmosphere of 20% CO_2_, 10% H_2_, and 70% N_2_ at 37°C. When necessary, antibiotics were added to the medium as follows: 7 µg/mL vancomycin, 100 µg/mL ampicillin, 200 µg/mL gentamicin, 20 µg/mL erythromycin for selecting *P. copri* DSM 18205 derived plasmid integrants; 5 µg/mL for other *P. copri* strains, and 20 µg/mL tetracycline for RHA03 and HDE06; 2.5 µg/mL for HDD04, HDE04 and HDD05.

##### Isolation of *P. copri* from humans

*P. copri* was isolated from fecal samples of *P. copri*-positive donors previously determined by 16S rRNA sequencing. Briefly, the fresh fecal samples were collected and further processed in an anaerobic chamber. A pea-sized fecal pellet was resuspended in 5 mL BHI+S and filtered through 70 µm cell strainer. We performed a serial ten-fold dilution of the flow-through, and streaked out the diluted samples with the dilution factors of 10^−3^ to 10^−6^ on BHI blood agar plates supplemented with vancomycin. The plates were then incubated anaerobically at 37°C for 48-72 hr. Individual colonies were picked into BHI+S broth and the resulting cultures were screened by PCR for *P. copri*-positive cultures using *P. copri*-specific primers (P_copri_69F/P_copri_853R). The pure *P. copri* isolates were obtained by steaking out the *P. copri*-positive cultures above, and confirmed by Sanger sequencing using the 16S rRNA gene-specific primers as described previously (16S_27F/16S_1492R) (Miller et al., 2013). The fresh culture of *P. copri* was mixed with an equal volume of 50% glycerol in BHI medium in sealed glass vials as bacterial glycerol stocks, and cryopreserved at −80°C immediately.

##### DNA extraction from human feces and *P. copri* cultures

The DNA extraction from fecal samples of *P. copri*-positive donors or *P. copri* strains cultured in BHI+S broth (OD_600_=0.6) was performed using ZymoBIOMICS DNA Miniprep Kit based on the instruction manual. We measured the concentration of purified DNA samples by Qubit Fluorometer (Thermo Scientific), and analyzed by agarose gel electrophoresis, NanoDrop^TM^ 2000 (Thermo Scientific), and Bioanalyzer (Agilent Technologies).

##### Whole genome sequencing, assembly, and annotation

The DNA library for genome sequencing of *P. copri* strains was performed using NEBNext® Ultra™ II FS DNA Library Prep Kit (New England Biolabs) for Illumina with parameters as followed: 500 ng input DNA and 5 min at 37°C for fragmentation; > 550-bp DNA fragments for size selection; primers from NEBNext Multiplex Oligos for Illumina Kit (New England Biolabs) for barcoding. The library was sequenced on the Illumina Miseq 2×250 bp The obtained reads were thus assembled with SPAdes version v3.10.0 using “careful” mode (Bankevich et al., 2012). Short contigs were then filtered by length and coverage (contigs > 500 bp and coverage > 5×). Gene prediction and annotation was performed using PROKKA version v1.13.3 (Seemann, 2014) with default parameters.

##### Phylogenetic analyses

Placement of *P. copri* complex. The phylogenomic analyses were conducted as previously described on the characterization of the *P. copri* Complex (Tett et al., 2019) using PhyloPhlAn3 (Asnicar et al., 2020) with reference set of *P. copri* strains (Tett et al., 2019) and the newly *P. copri* strains isolated in this study. The phylogenetic analysis in Figure 1A was built using the 400 universal marker genes of the PhyloPhlAn database using the parameters “--diversity low”, and “--accurate” option. The configuration file (config_file.cfg) was set with the following tools and parameters:

Diamond version v0.9.9.110 (Buchfink et al., 2015) with “Blastx” for the nucleotide-based mapping, “Blastp” for the amino-acid based mapping, and “--more-sensitive --id 50 --max-hsps 35 -k 0” in both cases. MAFFT version v7.310 (Katoh and Standley, 2013), with “--localpair -- maxiterate 1000 --anysymbol --auto” options. trimAl version 1.2rev59 (Capella-Gutiérrez et al., 2009), with “-gappyout” option. IQ-TREE multicore version v1.6.9 (Nguyen et al., 2015), with “-nt AUTO -m LG” options. RAxML version 8.1.15 (Stamatakis, 2014), with “-p 1989 -m GTRCAT -t” options.

For Figure 5 and S6, the amino acid sequences of HTCS, SusC-like, SusD-like, and SGBP-like proteins encoded by arabinan processing PULs (PUL^Ara^) from *Bacteroides thetaiotaomicron* VPI-5482, *Phocaeicola vulgatus* ATCC 8482 and 12 *P. copri* strains were used for the phylogenetic trees via the MEGA-X software, respectively. The evolutionary history was inferred using the Neighbor-Joining method (Saitou and Nei, 1987) with 1000 bootstrap replicates. The two types of PUL^Ara^ extracted from all recovered assemblies from the genus *Prevotella* (n=8), *Bacteroides* (n=13) and *Phocaeicola* (n=3) was calculated for the analysis of phylogenetic distributions, respectively.

##### Determination of *P. copri* sensitivity to oxygen

*B. thetaiotaomicron* VPI-5482, *P. copri* HDD04, HDB01, and DSM 18205 were grown in BHI+S broth anaerobically. The fresh bacterial cultures were divided into 1 mL aliquots into 2 mL tubes with the caps being open, respectively. These aliquots were aerobically incubated at 37°C. At four time points (0, 1, 2, 4 hrs), three aliquots from respective cultures were placed back to the anaerobic chamber and performed serial dilutions for counting CFUs.

##### Determination of *P. copri* sensitivity to antibiotics

The wells of a non-tissue culture flat bottom 96-well were loaded with 198 µl BHI+S media in the presence of 2-fold serial dilutions of erythromycin, tetracycline, chloramphenicol, spectinomycin, apramycin, and hygromycin ranging from 0.04 to 400 µg/mL. BHI+S media without antibiotics was loaded as a positive control. *P. copri* strains were grown in BHI+S broth to an optical density (OD_600_) of 0.5-0.7. 2 µL bacterial culture was then inoculated into each well (inoculation ratio of 1:100). Absorbance at OD_600_ of each well was measured at an interval of 1 hr for 5 days using the microplate reader (BioTek). Assays were performed in triplicate. To ensure that the concentrations of erythromycin and tetracycline used for selecting transconjugants of different *P. copri* strains were sufficient for killing all the wild-type *P. copri* cells, 1 mL of fresh *P. copri* culture (10^8^-10^9^ bacterial cells) was plated on the BHI blood agar with respective concentration of antibiotics in triplicate.

##### Prediction of HTCS genes in *P. copri*

The genes of *P. copri* HDD04 that encodes proteins containing all the domains of HTCS (PF07494-PF07495-PF00512-PF02518-PF00072-PF12833) according to the Pfam classification were identified as described previously (Terrapon et al., 2015). The hmmsearch was carried out using default parameters (Eddy, 1998)

##### Molecular cloning

The relevant primers and plasmids are described in Tables S1. PCR amplification for cloning was carried out using Q5 High Fidelity DNA Polymerase (New England Biolabs). The PCR products were purified, and followed by DNA assembling with PCR amplified plasmid using Gibson reaction (HiFi DNA Assembly Master Mix, New England Biolabs). The assembled products were transformed into *E. coli* β2155 by chemical transformation. The resulting colonies were randomly picked to detect the inserts and their sizes by colony PCR using OneTaq DNA Polymerase (New England Biolabs). Genetic modifications generated on plasmids were verified by sequencing at Microsynth Seqlab (Microsynth AG, Germany).

Specifically, the pEx-insertion vector was constructed as follows: Firstly, the thymidine kinase gene (*tdk*) and its promoter was deleted from the multiple cloning site using DNA assembling with PCR amplified plasmid, resulting in pExchange. Secondly, 300-bp of the *tuf* (elongation factor Tu, DSM18205_02600) promoter was inserted into pExchange exactly before the coding sequence of *ermG*, generating the pEx-insertion-ermG vector. For first trial of genetic insertion in *P. copri*, 3-kb DNA sequences from DSM18205_00642-43, 00941-42, and 02334-35 were cloned into the multiple cloning site of pEx-insertion-ermG as the homology arm for plasmid integration. The pEx-insertion-ermG with the DNA region from DSM_02334-35 (DSM_02334: putative β-glycoside hydrolase, *bgl*) was designated as pEx-insertion-ermG-DSM-bgl.

Based on pEx-insertion-ermG as the vector backbone, similar cloning procedures were performed for constructing various plasmids carrying: (1) different promoter sequences for driving selective marker; (2) different sizes of homology arms varying from 0.5-kb to 4-kb for integration in *P. copri* HDD04; (3) 3-kb cloned homologous regions from different *P. copri* strains in our collection; (4) a pLGB30-derived *tetQ* selective marker (García-Bayona and Comstock, 2019) instead of *ermG*; (5) the T1-T2 terminators copied from pSAM (Goodman et al., 2011) for blocking the transcriptional readthough for the HTCS genes after plasmid integration.

The pEx-deletion-ermG and pEx-deletion-tetQ vectors were created by inserting the counter-selection marker following *ermG* and *tetQ,* respectively. The counter-selection marker is a DNA fragment generated by splicing 300-bp of the *gdhA* (HDD04_01507) promoter and the *sacB* gene from the pEX18Ap plasmid (Hoang et al., 1998). The pEx-deletion-ermG-bgl was similarly generated as described above. For in-frame deletion of genes in *P. copri*, the approximately 2-kb regions flanking the target gene were amplified, and assembled with PCR amplified pEx-deletion-ermG. For gene complementation, the target gene flanking with approximately 1-kb up- and down-stream regions were entirely amplified, and cloned into pEx-deletion-ermG.

##### Genetic manipulations of *P. copri*

Overnight culture of the *E. coli* donor strain was subcultured into LB medium containing ampicillin and DAP and *P. copri* subcultured into BHI+S medium. When they were grown to exponential phase (OD_600_=0.5-0.7), *E. coli* culture was transferred into the anaerobic chamber. The following procedures of genetic manipulations for *P. copri* including plasmid insertion, in-frame deletion, and complementation were performed in the anaerobic chamber. For conjugation, 1 mL *E. coli* culture (∼5×10^9^ CFUs) was centrifuged at 8000× g for 3 min to pellet the bacterial cells, followed by resuspension in 100 µL fresh *P. copri* culture (∼5×10^7^ CFUs) to get a ratio of donor: recipient of 100: 1. Specially, if *P. copri* HDA04 or HDD12 culture was used as the recipient strain for conjugation, to obtain the same donor/recipient ratio above, 20 µL HDA04 culture plus 80 µL BHI+S medium or 10 µL HDD12 culture plus 90 µL BHI+S medium was used to resuspend the *E. coli* pellet, respectively. The resuspension was then plated on a BHI blood agar with DAP for 18 hrs at 37°C for bacterial conjugation unless otherwise stated. Bacterial cells were washed off from the plate using 1 mL BHI+S medium, mixed well, and plated serial dilutions or the whole bacterial pellet after centrifugation on BHI blood agar plates containing gentamicin in addition of erythromycin or tetracycline. Colonies generated from transconjugants were visible after incubation of plates for 2-4 days according to properties of the *P. copri* derivatives. If necessary, the CFUs were counted for quantification of transconjugant yields. Insertion of the plasmid was verified by amplifying two joints between the bacterial chromosome and vector via colony PCR using P3/P4 and P5/P6 primer pairs, with P1/P2 amplified DNA as a control.

For in-frame deletion and complementation, the insertion mutants were grown in liquid BHI+S without selection, and then subcultured every 12 hr for allelic exchange. The final culture was plated onto YT agar plates supplemented with 5% sucrose to select the revertants (wild type) and gene deletion mutants with loss of the vector. After incubation of plates for 2-4 days, individual colonies were restreaked onto BHI blood plates in the presence and absence of erythromycin using the same inoculating loop, respectively, to further confirm erythromycin sensitivity of the clones. Erythromycin-sensitive clones were subsequently screened for the genetic modifications (gene deletion or complementation) by PCR and verified by sequencing at Microsynth Seqlab (Germany).

##### Prediction of PULs in *P. copri* genomes

The prediction of PULs in *P. copri* genomes and MAGs was described previously (Gálvez et al., 2020). Briefly, PULs and *susC/D*-like gene annotations were carried out using PULpy (Stewart et al., 2018) (commit 8955cdb, https://github.com/WatsonLab/PULpy). Annotation of carbohydrate-active enzymes (CAZymes) surrounding the *susC*/*susD*-like pairs was performed by using dbCAN2 tool (Zhang et al., 2018) version v2.0.6 (CAZy-DB=07312019, https://github.com/linnabrown/run_dbcan).

##### Measurement of *P. copri* growth on a carbohydrate array

The growth curves of *P. copri* strains cultured in minimal medium (MM) supplemented with a sole carbohydrate were measured as previously described with the following modifications (Martens et al., 2011). The wells of a non-tissue culture flat bottom 96-well were loaded with 100 µl sterilized carbohydrate stocks (2× concentration). Each carbohydrate was added into at least three wells. *P. copri* was grown in MM+Glucose to an OD_600_ value of approximately 0.6. 400 µL culture was then centrifuged to pellet the bacterial cells. The pellet was washed by 1 mL 2× MM without any carbohydrates and resuspended in 10 mL 2× MM as a seed culture. Each well of the plate was loaded with 100 µL seed culture. Absorbance at OD_600_ of each well was measured for 5 days by the microplate reader (BioTek) at 1-hr intervals with 15-second pre-shaking.

In Figure 2D, Figure S5, and Table S3, the maximal OD_600_ values subtracting the background reads (OD_600_ max) in the curves were identified for calculating means and standard deviations (SDs). Because we observed that the presence of erythromycin in MM significantly affect the duration of lag phase, but not the growth pattern of *P. copri*. The growth curves of HTCS gene insertion mutants and relevant intergenic insertion control were therefore shown starting from the OD_600_ values increased by 10% of the OD_600_ max to the OD_600_ max in Figure 2B and 2C.

##### RNA extraction from human feces and metatranscriptome sequencing

The fecal sample from the human donor carrying *P. copri* HDD04 was immediately collected into DNA/RNA Shield Fecal Collections Tubes and stored at 4°C for stabilizing RNA. An aliquot of 400 µL content from the tube was used for isolating RNA using ZymoBIOMICS RNA Miniprep Kit following the instruction manual.

##### RNA extraction and RNA-seq library preparation

*P. copri* HDD04 was grown in BHI+S broth to the exponential phase (OD_600_=0.6). 5 mL fresh cultures were treated by RNAprotect (New England Biolabs) based on the manufacturer’s instructions, pelleted by centrifugation, and stored at −80°C until further processing. The bacterial RNA was isolated using ZymoBIOMICS RNA Miniprep Kit following the instruction manual. RNA quality was evaluated by agarose gel electrophoresis, NanoDrop^TM^ 2000 (Thermo Scientific), and Bioanalyzer (Agilent Technologies) according to RNA integrity score (RIN > 8.0). Bacterial ribosomal RNA (rRNA) was then depleted by Ribo-Zero Gold rRNA Removal Kit (Epidemiology) as described in the commercial protocol. Libraries for Illumina sequencing were prepared using the NEBNext® Ultra™ Directional RNA Library Prep Kit for Illumina® (New England Biolabs) following manufacturer’s protocol. For each sample, 100 ng of fragmented mRNA was used as an input for cDNA synthesis and Illumina sequencing adaptor ligation.

For other treatment in Figure 4, *P. copri* HDD04 was initially grown in minimal media plus glucose (MM+Glucose) to an OD_600_ value of 0.5. 2 mL culture was then centrifuged to pellet the bacterial cells and resuspended using an equal volume of minimal media without carbohydrates (MM), followed by another centrifugation and resuspending in the same volume of MM. 40 µL suspension was inoculated into 4 mL MM plus glucose, arabinan, arabinoxylan, pectic galactan, and inulin, respectively. Three replicates were performed for each carbon source. Once *P. copri* grown to OD_600_=0.5, 750 µL culture was taken and treated by RNAprotect (New England Biolabs). Bacterial ribosomal RNA (rRNA) was thus depleted using Pan-prokaryote riboPOOL™ Kit (siTOOLs Biotech) as described in the manual. The cDNA library preparation and sequencing was carried out as described above.

##### RNA-seq analysis

Reads were quality filtered using Trimmomatic (Bolger et al., 2014) version v0.33 with as follow parameters (LEADING:3 TRAILING:3 SLIDINGWINDOW:4:15 MINLEN:35 HEADCROP:3). After quality control reads were aligned to each *P. copri* reference genome using STAR (Dobin et al., 2013) version v2.5.2a. Reads count was performed using HTSeq (Anders et al., 2015) version v0.11.2. With the aim to control for interspecies multi-mapping, reads were split by mapping to multiple references using (BBsplit). References genomes were selected from the reconstructed MAGs and one representative strain for each of the *P. copri* clades in combination with each donor’s isolate from the clade A.

For *in vivo* and *in vitro* differential gene expression, gene read counts were transformed using TPMs normalization and differential gene expression was quantified in R using the DEseq analysis with a single replicate (iDEG) package (Li et al., 2019).

For the transcriptome in vitro with supplemented polysaccharides, samples were proceeded as described above. Normalization and differential expression were quantified in R using the DEseq2 package (Love et al., 2014) version v1.26.0 using the samples grown in MM + glucose as a control.

##### Measurements of gene transcription by RT-qPCR

The preparation of *P. copri* cultures and extraction of total RNA were performed as described above. Reverse transcription was carried out with ProtoScript® II First Strand cDNA Synthesis Kit (New England Biolabs) using Random Primer Mix and 800 ng purified RNA as template for 20 µL reaction. The abundance of transcript for target *susC*-like genes and reference *tuf* gene was quantified with KAPA SYBR® FAST qPCR mix (KAPA Biosystems) using 0.5 ng/µL template cDNA, 25 nM of each target gene-specific primer. The reaction was performed in a 96-well plate on the Roche Lightcycler 480. Using the ddCT method, raw values were normalized to values for the *tuf* gene and then the fold change was calculated by dividing MM+specific polysaccharide values by values obtained from MM+glucose.

##### Reconstruction of *P. copri* MAGs

The reconstruction of *P. copri* MAGs from a recent dataset (De Filippis et al., 2019) was performed as described previously (Gálvez et al., 2020). In brief, the sequencing data of the gut microbiome from 101 healthy Italian individuals with distinct diets (Omnivore, n = 25; Vegetarian, n = 39; Vegan, n = 37; NCBI SRA: SRP126540 and SRP083099) was analyzed as follows: (1) Sample-wise assembly, annotation, and integrative genomic binning was carried out with ATLAS metagenomic workflow (Kieser et al., 2020) (commit a007857, https://github.com/metagenome-atlas/atlas); (2) Genome abundance estimates were calculated for each sample by mapping the reads to the non-redundant MAGs using BBmap and determining the median coverage across each of the MAGs.

### QUANTIFICATION AND STATISTICAL ANALYSIS

#### Statistical Analysis

Statistical analyses were carried out in R (R Core Team, 2019) and figures were produced using 690 the package ggplot2 (Wickham, 2016). Datasets were analyzed using the GraphPad Prism 8. Pairwise comparisons were performed using Student’s *t* test with a paired, two-tailed distribution. More statistical details are indicated in the associated figure legends when required.

## DATA AND SOFTWARE AVAILABILITY

The accession numbers for all whole genome sequencing and 16S rRNA data reported in this manuscript are available under NCBI BioProject ID: PRJNA684333.

Transcriptome analysis and R customised code is available in http://github.com/strowig-lab/galvez_et_al_2020/.

## ACKNOWLEDGMENTS

We thank members of the Strowig laboratory for valuable discussions. We thank Andrew Goodman for helpful discussions. We thank the genome analytics core facility of the Helmholtz Institute for Infection Research. J. L. was funded by Singh-Chhatwal Stipend from Helmholtz Centre for Infection Research and Humboldt Research Fellowship from Alexander von Humboldt Foundation.

## AUTHOR CONTRIBUTIONS

J.L. and T.S. designed the experiments and wrote the paper. J.L. conducted the most of experiments and data analysis. L.A. and A.I. performed the strain isolation. J.L., E.J.C.G., and T.R.L performed the bioinformatic analysis. L.A. and E.A. assisted in grow assays. L.A. and A.A.B assisted in RNA preparation for RNA-seq. E.A. assisted in molecular cloning and genetic manipulation.

